# The effects of chronic consumption of lipid-rich and delipidated bovine dairy milk on brown adipose tissue volume in wild-type mice

**DOI:** 10.1101/2021.11.07.467371

**Authors:** Zachary D’Alonzo, John Mamo, Liam Graneri, Ryu Takechi, Virginie Lam

**Affiliations:** Curtin Health Innovation Research Institute, Curtin University, Perth, WA 6845, Australia; Curtin Medical School, Faculty of Health Sciences, Curtin University, Perth, WA 6845, Australia; School of Population health, Faculty of Health Sciences, Curtin University, Perth, WA 6845, Australia

**Keywords:** Milk, hypercaloric diet, brown adipose tissue, body composition

## Abstract

Brown adipose tissue (BAT) activation is associated with increased energy expenditure by inducing non-shivering thermogenesis. Ingestion of a milk fat globule membrane (MFGM) supplement and a high calorie diet are reported gateways into BAT activation. However, little is known about the effect of MFGM and high calorie diets on BAT volume. To gain insight into this, mice were maintained on a high fat (HF) or low-fat (LF) diet in conjunction with either full-cream (FC) or skim bovine dairy milk (BDM). After being maintained on their respective diets for 13 weeks, body composition, including BAT volume, was measured using X-ray microtomography. A high calorie diet resulted in an increase in BAT volume and mice consuming a HF diet in conjunction with FC BDM had significantly greater BAT volume than all other groups. Conversely, mice consuming a HF diet in addition to skim milk had lower BAT volume compared to the HF control. The data presented suggests that consumption of a high calorie diet in conjunction with FC BDM increases BAT volume in wild-type mice. This study may provide valuable insight into future studies investigating BAT volume and BAT activity in relation to environmental factors including diet.

## 1. Introduction

Mammals regulate heat production through two mechanisms known as shivering and non-shivering thermogenesis (NST) (1). Shivering thermogenesis defines the conversion of chemical energy into heat through generation of a contraction in specific muscle groups in response to a cold stimulus. On the contrary, NST defines the biochemical process of heat generation which does not induce shivering and is considered a more efficient method of thermoregulation. Brown adipose tissue (BAT) is considered the primary site of NST, where NST takes place by inducing rapid intracellular lipolysis by activation of adipose triglyceride lipase and hormone sensitive lipase (2–4). This ultimately results in the release of fatty acids by lipid droplets which thereafter can be converted into heat via uncoupling protein 1 (UCP1) in mitochondria (5, 6). Conversely, white adipose tissue (WAT) has the opposite role and stores excessive energy as triglycerides (7).

Historically, BAT had been thought to have little significance in adult humans (7). However, within the past two decades, a number of studies have reported that higher BAT mass is associated with lower BMI (8, 9). Furthermore, the activation of BAT is increasingly recognised as an important process for BAT’s energy utilisation. Whilst non-activated BAT has metabolic activity comparable to that of WAT (10), activated BAT has increased thermogenic capabilities, consequentially resulting in increased energy expenditure and may therefore contribute to fat loss (11). Additionally, evidence suggests that activated BAT has beneficial metabolic effects and reportedly regulates blood glucose levels in type 2 diabetes patients (12).

BAT can be activated by cold exposure or induced by ingestion of a meal (diet-induced thermogenesis (DIT)). Recent evidence suggests that the milk fat globule membrane (MFGM), the membrane that encompasses milk fat and allows for efficient distribution of milk lipids, can attenuate body weight gain via the activation of BAT in mice maintained on a high-fat diet (13). Mice that were fed high-fat chow supplemented with an infant MFGM had reduced weight gain of up to 34.82%, which was positively associated with the quantity of MFGM consumption compared to the mice fed only with high-fat chow (13). Treatment of mice with MFGM led to an upregulation of UCP1 expression in BAT, indicative of significantly enhanced conversion of triglycerides into heat. Moreover, MFGM and phosphatidylcholine were shown to accelerate the conversion of white to beige adipocytes that have thermoregulatory abilities comparable to BAT (14).

However, no studies to date have investigated the effects of commercially available bovine dairy milk (BDM) on BAT volume and weight. Thus, in the present study we evaluated the effect of full cream BDM on BAT and WAT abundance in comparison with delipidated, skim BDM in C57BL/6J mice, and how BDM fat can affect the production of BAT and WAT in mice maintained concurrently on a high or low-fat diet.

## 2. Materials and Methods

### 2.1 Animals and dietary intervention

Young adult wild-type C57BL/6J male mice were purchased from Animal Resources Centre (WA, Australia) at 5 weeks of age. Following a 1-week acclimatisation period, mice were randomly separated into one of 5 dietary intervention groups (n=5 per group). The low-fat control group was given water with a standard maintenance chow (LF) (AIN-93M, Specialty Feeds, WA, Australia). The high-fat control group was given water with a high-fat chow (HF) (SF07, Specialty Feeds, WA, Australia). Two additional groups were maintained on 20% full cream milk solution diluted in water with one group given the low-fat diet (LF+FC) and one group receiving the high-fat chow (HF+FC). The final group was given 20% skim milk solution diluted in water in addition to the high-fat chow (HF+Skim). Each group was sacrificed 13 weeks after commencing the dietary intervention. Mice were housed in groups of 5 in ventilated cages with a 12h light/dark cycle under controlled air pressure and temperature (2ºC). All groups had ad libitum access to food and drinks. This study was approved by the Curtin Animal Ethics Committee (AEC Approval No. 2018-03).

### 2.2 In vivo body fat composition analysis

After 13 weeks of dietary intervention, body composition was measured using Skyscan high resolution in-vivo X-ray microtomography (Bruker, MA, USA) at the Centre for Microscopy, Characterisation and Analysis (Harry Perkins Institute, WA, Australia) as previously described (15). Briefly, mice were anesthetised with isofluorane gas, and the thoracic and abdominal areas of each mouse was imaged with 40kV and 383 μA intensity. To limit noise and artefacts, smoothing was set to 2 and ring artefact reduction was set to 6. To optimise contrast, beam hardening correction was set to 20%. Total body fat percentage, subscapular BAT and total body WAT volumes were calculated through tomography intensity grading of 3-D X-ray shadow projection reconstructions using 3-D rendering software CT-Analyser (Bruker).

### 2.3 Statistical analysis

All data is displayed as mean ± SEM. All data were analysed with one-way analysis of variance (ANOVA) followed by Fisher’s least significant difference (LSD) post hoc test for multiple comparison (GraphPad Prism 9, USA). Statistical significance was detected at p < 0.05.

## 3. Results

### 3.1 Food and liquid consumption

Average daily liquid and food intake per mouse was measured and is presented in figure 1. Food and liquid intake were used to determine total cumulative energy from macronutrients lipids, carbohydrates and proteins over the 13-week period. LF and HF control groups consumed similar liquid quantities and significantly less compared to all milk groups. HF+FC and HF+Skim groups consumed similar liquid quantities which were slightly higher than the control groups. The LF+FC group consumed the greatest quantity of liquid and was significantly higher than all the other groups. Food consumption was similar across all groups with only the HF groups being slightly but significantly higher than the LF+FC group.

**Figure 1.**
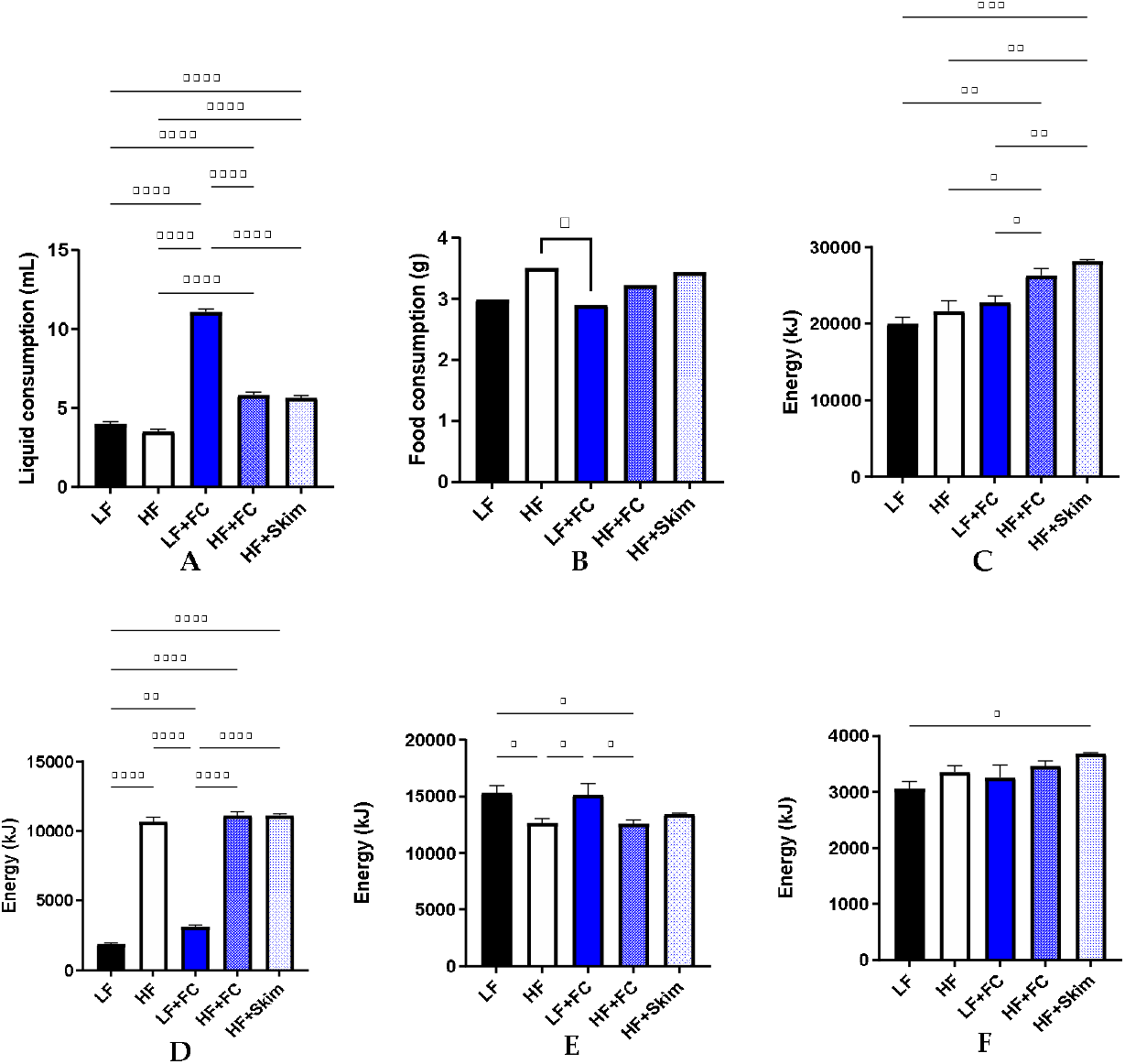
Liquid consumption (A) and food consumption (B) was measured daily per mouse over the 13-week interventions. Total energy consumption (C), total energy from lipids (D), total energy from carbohydrates (E) and total energy from protein (F) was also measured following 13-week interventions. Energy consumptions are cumulative following the 13-week intervention.

Both HF+Skim and HF+FC groups had significantly higher total energy intake than HF group and LF groups, with HF+Skim group consuming the most energy, followed by HF+FC. All HF groups had similar and significantly higher cumulative energy intake from lipids as a result of the ingestion of high-fat diet. LF and LF+FC had the highest cumulative energy intake from carbohydrates, which were significantly higher than all HF groups except HF+Skim. All groups had similar energy intake from protein as all food and milk solutions contained adequate amounts of protein. However, HF+Skim maintained a slightly higher level of protein intake than the other groups which was significantly higher than the LF group.

### 3.2 Dietary intervention effect on body composition

Mass and body composition of the mice are depicted in figure 2. All HF groups had significantly higher body mass compared to all the other LF groups. Body WAT volume in LF was the lowest, followed by LF+FC, both significantly lower than all the other groups receiving HF. Despite HF+Skim and HF+FC having similar body mass, HF+FC had significantly higher body WAT volume than HF+Skim. The HF had the lowest body mass out of the HF groups, yet significantly higher than LF or LF+FC. The consumption of HF alone or FC milk alone did not significantly increase the BAT volume compared to the mice maintained on LF. However, the combination of HF+FC significantly increased the BAT volume compared to all other groups. The synergistic effects of HF and FC milk intake was not realised when the mice received HF and Skim milk. The ratio of BAT volume per gram of body weight was used to clearly describe BAT measured relative to the size of an individual mouse (Fig 2D). Relative to the total body mass, the mice maintained on HF+FC showed the significantly higher BAT volume compared to all the other groups.

**Figure 2.**
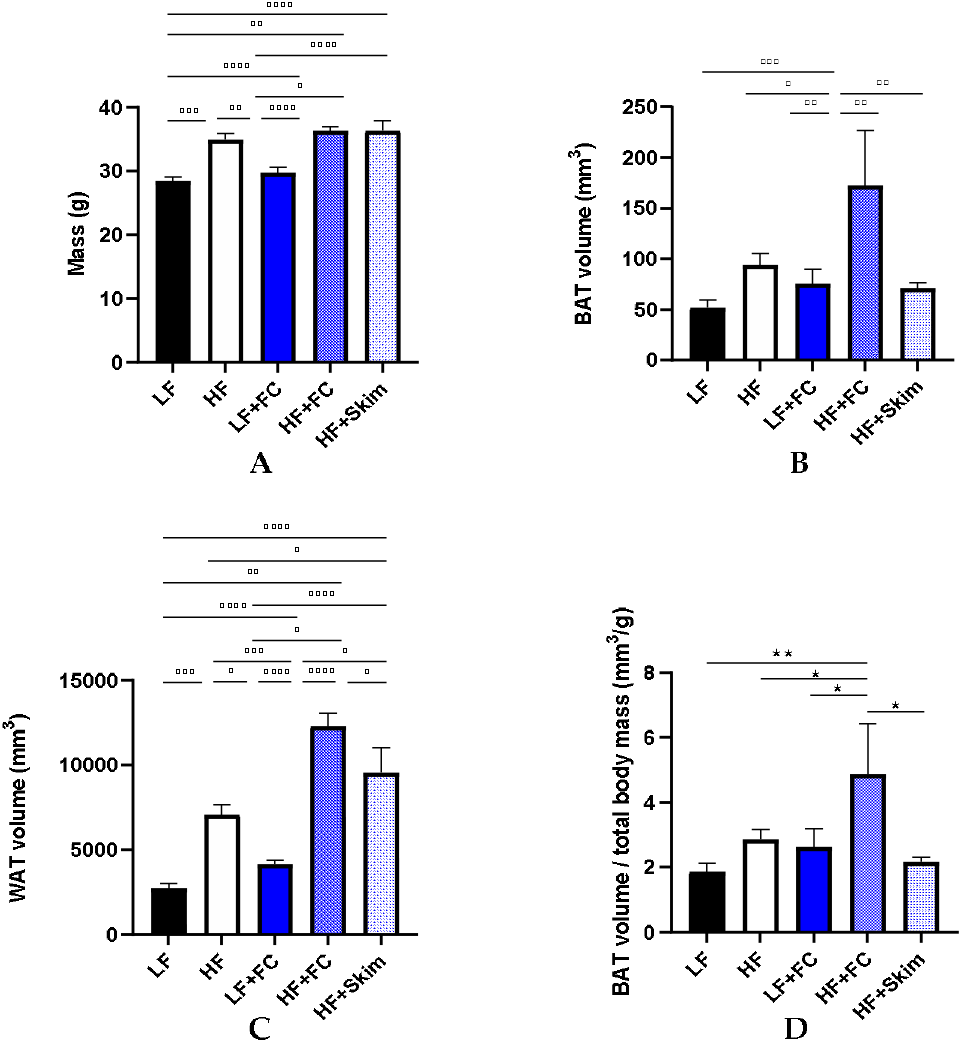
Mice body mass (A) and body composition analysis showing in-vivo X-ray microtomography results of BAT volume (B) and WAT volume (C). BAT volume per gram of body weight (D) was also measured.

## 4. Discussion

BAT is an increasingly researched organ that has known roles in NST and until recently, was thought to be physiologically irrelevant in the adult human. However, recent research has shown that environmental factors such as cold temperature and certain diets can influence adult BAT activity. Therefore, in this study we report the effects of the chronic intake of BDM and hypercaloric HF diet on BAT volume in young adult wild-type mice.

Intervention with a diet enriched in fat (HF) for 13 weeks almost doubled the BAT volume in mice compared to the mice receiving low-fat control chow (LF). Similar observations were reported by previous clinical and pre-clinical studies, which identified a significant increase in BAT activity after chronic hypercaloric feeding and even after a single hypercaloric meal, (16–18). However, these studies did not determine the volume of BAT. Thus, our study for the first time demonstrates that high calorie HF diets increase BAT volume.

Mice receiving a HF diet in conjunction with full cream bovine dairy milk (HF+FC) displayed significantly higher net BAT volume as well as BAT proportion per total body mass, compared to every other group in the study. Despite consuming an almost identical amount of calorie and macronutrients over the study period, the mice on HF diet in conjunction with delipidated skim milk (HF+Skim) group showed significantly lower BAT volume compared to the HF+FC mice. Similarly, the mice that consumed only FC BDM without hypercaloric HF diet (LF+FC group) showed significantly less BAT volume compared to the mice in HF+FC group. The combination of FC BDM and the HF diet not only introduces additional lipids to a lipid-rich diet, but also associated with an influx of proteins and phospholipids that form the MFGM. Although we did not analyse BAT activation following the respective interventions, it is known that MFGM supplementation in mice increases activity of BAT (13, 14). This study suggests that the ingestion of FC BDM in conjunction with a high-fat diet may result in increased MFGM, which may consequently increase BAT volume.

Interestingly, both FC and skim milk in combination with HF diet showed significant increase in WAT volume, whilst only mice with HF+FC showed the BAT volume increase. This suggests that the intake of FC milk but not skim milk with HF diet may promote the browning of WAT. Irrespective of WAT volume, studies suggest that certain nutritional interventions and exercise regimes may promote browning of WAT to produce beige adipocytes to modulate energy expenditure (19–21). These studies also suggest that the browning of WAT typically goes coincides with increases in BAT activity. Of particular importance, Li et al established that MFGM may induce browning of WAT alongside BAT activation (13, 14). Therefore, the increase of BAT volume in HF+FC mice suggests that browning may be taking place in the WAT of this intervention group.

## 5. Conclusions

In this study we have shown that a high-fat, hypercaloric diet together with FC BDM increases BAT volume in wild-type mice. Further intervention studies with BDM will be required in order to identify the full impact on BAT activity, WAT browning and energy expenditure in human subjects.

## Supplementary materials

The following are available online at www.mdpi.com/xxx/s1, Figure S1: Macronutrient composition of dietary intervention components

**Supplementary Table 1.**
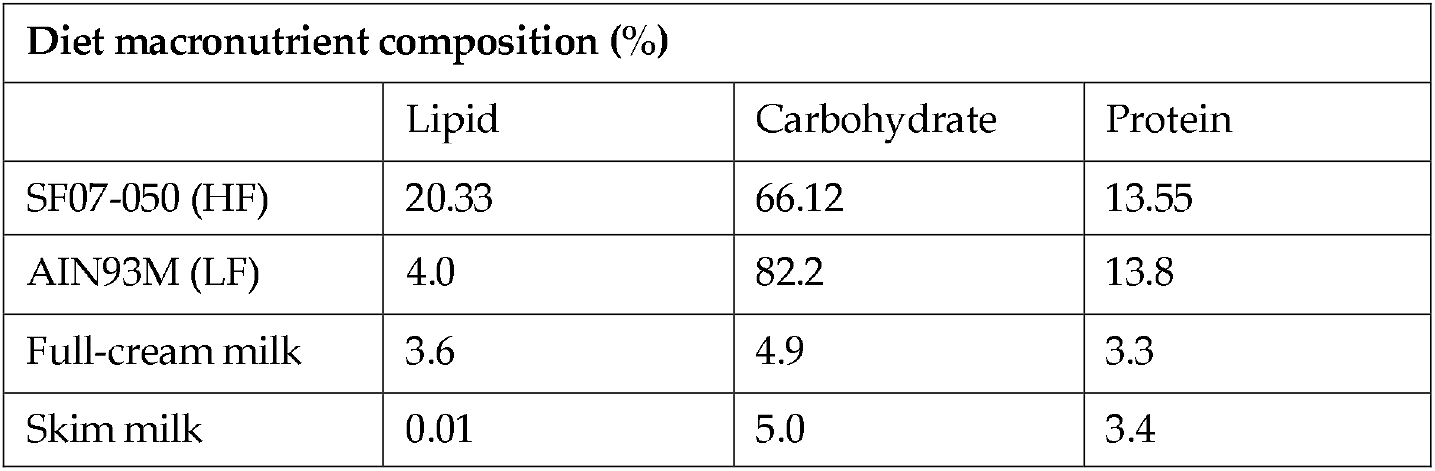
Macronutrient composition of dietary intervention components

| Diet macronutrient composition (%) |  |  |  |
| --- | --- | --- | --- |
|  | Lipid | Carbohydrate | Protein |
| SF07-050 (HF) | 20.33 | 66.12 | 13.55 |
| AIN93M (LF) | 4.0 | 82.2 | 13.8 |
| Full-cream milk | 3.6 | 4.9 | 3.3 |
| Skim milk | 0.01 | 5.0 | 3.4 |

**Supplementary Figure 1.**
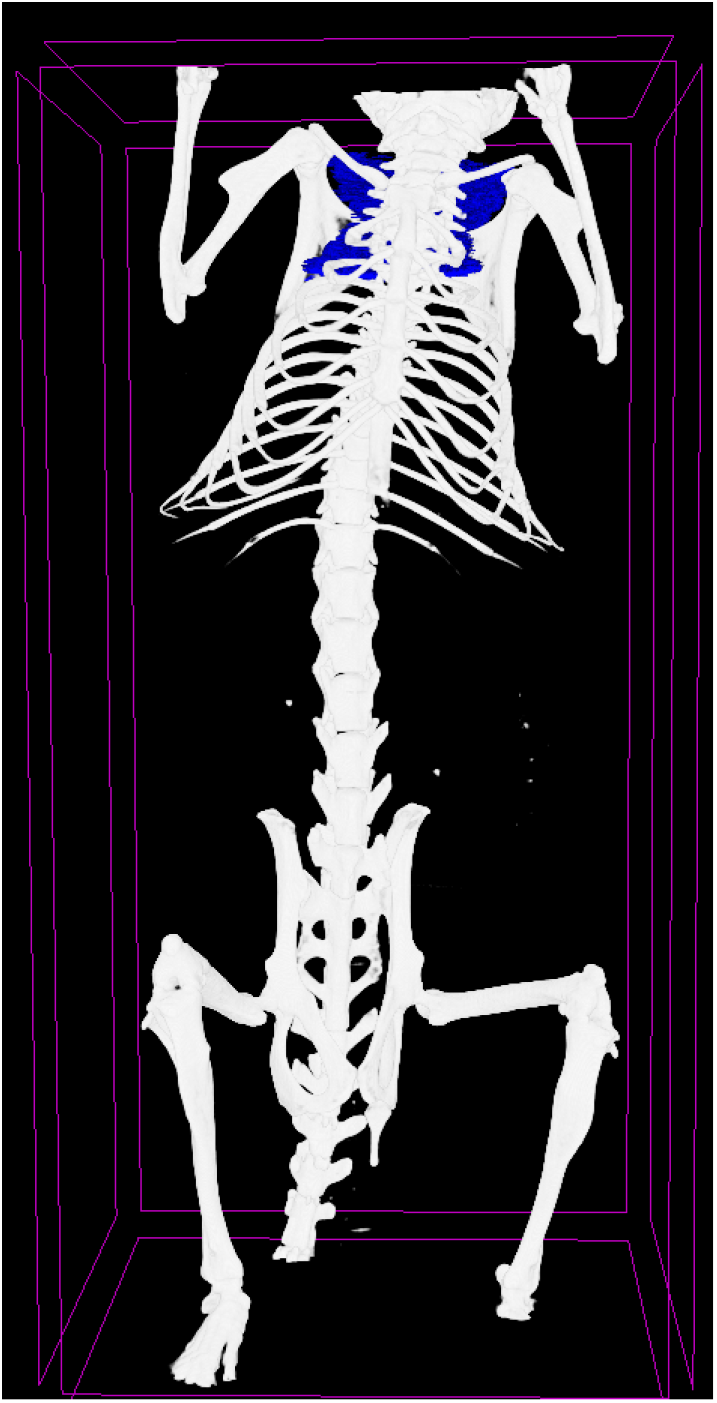
Representative SkyScan image of mouse subscapular BAT (blue) from HF+FC group.

## Author Contributions

Conceptualisation, Z.D., J.M., R.T. and V.L.; methodology, Z.D. and L.G.; software, Z.D. and L.G.; validation, Z.D., J.M., L.G., R.T. and V.L.; formal analysis, Z.D.; investigation, Z.D., R.T. and V.L.; resources, J.M., R.T. and V.L.; data curation, Z.D. and R.T.; writing-original draft preparation, Z.D. and R.T.; writing—review and editing, Z.D., R.T. and V.L.; visualization, Z.D., R.T. and V.L.; supervision, J.M., R.T. and V.L.; project administration, J.M., R.T. and V.L.; funding acquisition, J.M., R.T. and V.L. All authors have read and agreed to the published version of the manuscript.

## Institutional Review Board Statement

The study was conducted according to the guidelines of the Declaration of Helsinki, and approved by the Curtin Animal Ethics Committee (protocol code 2018-03, 2018)

## Data Availability Statement

All data is presented and available in this manuscript.

## Acknowledgments

The authors acknowledge CMCA and the technical staff, Diana Patalwala and Ivan Lozic, for their assistance in Skyscan analyses.

